# An anesthetized rat assay for evaluating the effects of organophosphate-based compounds and countermeasures on intracranial EEG and respiratory rate

**DOI:** 10.1101/2024.05.15.594373

**Authors:** J.S. Thinschmidt, J.D. Talton, S.W. Harden, C.J. Frazier

## Abstract

The development of medical countermeasures (MCMs) against organophosphate (OP) induced poisoning is of substantial importance. Use of conventional therapeutics is complicated by off-target effects and restricted penetration of the blood-brain barrier (BBB). Therefore, a concerted effort is underway to discover improved acetylcholinesterase (AChE) reactivators, muscarinic acetylcholine receptor (mAChR) antagonists, and other countermeasures with broader spectrum activity and enhanced CNS efficacy. We recently developed a rat brain slice assay to assess the efficacy of AChE reactivators and mAChR antagonists against the acute effects of the organophosphorus AChE inhibitor 4-nitrophenyl isopropyl methylphosphonate (NIMP) in the basolateral amygdala (BLA). Here we introduce a complimentary anesthetized animal model to evaluate the same compounds *in vivo* with concurrent monitoring of EEG and respiratory rate. We find that intravenous delivery of 0.5 mg/kg NIMP reliably produces seizure like activity in the BLA, with concurrent respiratory depression and eventual respiratory failure. The central effects of AChE reactivators and mAChR antagonists delivered intravenously are consistent with their expected ability to cross the BBB. Combining our previously described *in vitro* assay with the methods described here provides a relatively comprehensive set of preclinical tools for evaluating the efficacy of novel MCMs. Notably, using these methods potentially obviates subjecting conscious animals to cholinergic crises, which aligns with the AAALAC’s 3Rs principle of refinement.

## 1 Introduction

Organophosphates (OPs) were extensively utilized in mid-20^th^ century pesticides. They continue to be employed in that role, and are also used as chemical weapons. These compounds inhibit acetylcholinesterase (AChE) by phosphorylating its active site, preventing the breakdown of acetylcholine (ACh) [1,2] (Figure 1). This inhibition leads to detrimental effects on both the peripheral and central nervous systems [3,4]. In the central nervous system, the acute effects of OP-based toxins are often fatal, manifesting as loss of consciousness, seizures, and potential long-term brain damage due to excitotoxicity. Standard first-line treatments since the 1970’s for OP-based nerve agent exposure include atropine sulfate (a nonselective muscarinic acetylcholine receptor antagonist) in combination with the AChE reactivators pralidoxime chloride (2-PAM). However, AChE reactivators are reported in the literature to exhibit limited penetration of the blood-brain barrier (BBB) and therefore to have limited effectiveness against central effects of OPs [e.g. see 5,6]. This underscores the need for the development of novel AChE reactivators capable of more effectively penetrating the BBB to counter OP-induced CNS effects. The ideal next generation reactivators should be able to effectively restore AChE inhibited by multiple OPs. Also, the off-target actions of atropine can produce unwanted side effects, emphasizing the requirement for compounds designed to be highly selective for receptors involved directly in OP-induced hyperexcitation in the CNS.

**Figure 1:**
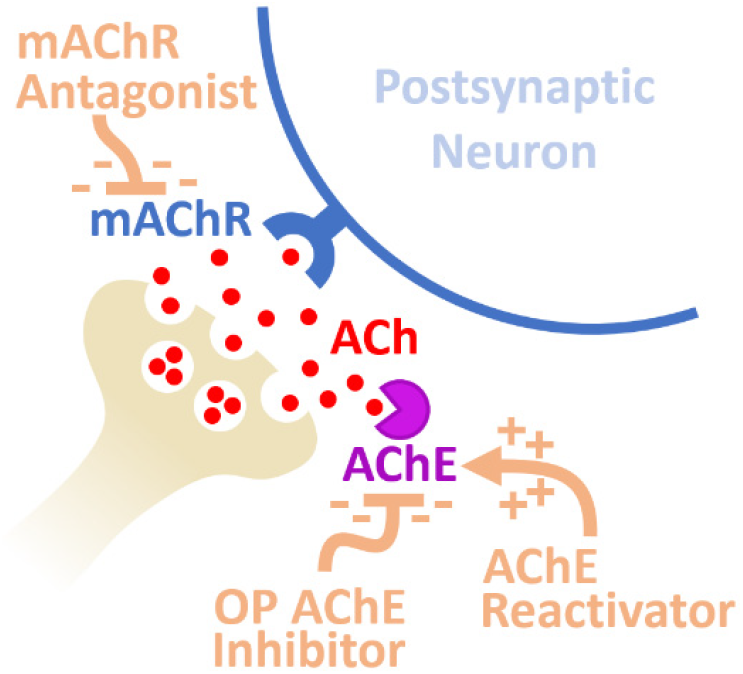
Diagram of synaptic mechanisms mediating neuronal hyperexcitation following exposure to OP-based AChE inhibitors. Inhibition of AChE results in a substantial increase in mAChR activation from excessive ACh at the synapse producing hyperexcitation. Medical countermeasures against OP poisoning include AChE reactivators and mAChR antagonists.

Developing novel therapeutics requires preclinical investigations utilizing models that are relevant for the desired clinical outcomes. In developing medical counter measures (MCMs) for OP-poisoning in the CNS, animal models should aim to evaluate seizure prevention and cessation along with prevention of long-term neurological consequences. The validity of such work regarding its clinical translation to humans is worth consideration. Acute toxicity from OPs in diverse animal species, including rodents and large mammals, closely mirrors human exposure symptoms. High doses of OPs induce a cholinergic crisis in conscious animals, marked by manifestations such as chewing, gnawing, profuse secretions, diarrhea, muscle fasciculation, restlessness, tremors, convulsions, and fatal respiratory distress [3,7–10]. The correlation between LD50s and *in vitro* acetylcholinesterase (AChE) inhibition in brain extracts underscore AChE inhibition as a critical factor in OP-induced toxicity in animal models [3,8,11–14]. Further, the correlation between lethality following OP exposures and the presence of seizures has been well documented in rodents [15], and is consistent with the increased risk of death associated with status epilepticus in humans [16,17]. In summary, the signs of acute OP poisoning and long-term CNS effects in animals closely resemble clinical outcomes in humans.

We recently developed an *in vitro* assay that enabled quantification of the effects of acute exposure to OPs and MCMs in the brain [18]. We used rat brain slices containing the basolateral amygdala (BLA) because the BLA receives dense cholinergic innervation [19], expresses high levels of AChE [20,21], and contributes directly to status epilepticus following exposure to soman [22,23]. We found the excitatory effects of bath-applied ACh on voltage-clamped BLA pyramidal neurons and excitatory synaptic transmission were strongly enhanced by the OP-based AChE inhibitor NIMP. Further, we demonstrated these effects were dependent on activation of M1 mAChRs and could be reversed with the AChE reactivator HI-6. Overall, the study outlined a novel approach to quantifying the effects of OP-based AChE inhibitors and reactivators in the CNS, and strongly reinforced a role for the BLA in producing CNS hyperactivity resulting from acute OP-poisoning.

While our work using brain slices provided valuable insights for MCM development and testing, it did not allow for simultaneous evaluation of peripheral effects and did not provide any information on BBB permeability. For these reasons, we developed a whole animal model to record both BLA EEG activity and respiratory function following acute OP poisoning and administration of MCMs. The assay is novel as recordings are acquired from isoflurane-anesthetized rats using IV administration of NIMP and MCMs, enabling the ability to monitor CNS hyperexcitation induced by OP actions on endogenous cholinergic activity. Included in the assay are intact cholinergic projection pathways and the ability to monitor respiratory rate, providing an assessment of OP-induced pathologies and the ability of novel and conventional MCMs to prevent or reverse them in whole animals.

Our previous work in brain slices and the work presented here outline two compassionate alternatives to testing in conscious animals during the drug discovery process for MCMs against OP-poisoning. In that regard, this study aligns well with the principle of “refinement” outlined by the Association for Assessment and Accreditation of Laboratory Animal Care (AAALAC). We suggest that the *in vivo* assay outlined in the current study, employed in combination with an in vitro assay recently described [18], offers an effective and comprehensive alternative to unanesthetized animal models for preclinical assessment of novel compounds as MCMs against OP-induced CNS hyperactivity.

## 2 Materials and Methods

### 2.1 Animals

Male Sprague Dawley rats (175-300g, 2-3 months old, Envigo, Indianapolis, IN) were housed in static cages. Standard rodent chow and water was available ad libitum. Housing was temperature and humidity controlled (20° - 26°C, and 30% - 70%, respectively). All procedures performed on live animals as described below were reviewed and approved by the Institutional Animal Care and Use Committee at the University of Florida.

### 2.2 Jugular vein catheterization

Procedures for jugular vein catheterization closely followed those outlined by Feng et al. [24]. Animals were placed in a plexiglass induction chamber with 3% isoflurane vaporized in 100% O_2_ for ∼5min. Following no withdrawal reflex, they were moved to a water-jacketed heating pad that maintained body temperature at 37°C and a nose cone was secured with tape to ensure continuous delivery of anesthesia. A 4-5 cm ventral cervical skin incision was made to the right of the neck midline at the level of the clavicle. Hemostats were used to bluntly dissect the right jugular vein exposing a 5-6 mm section. A loose tie was made with 4-0 silk suture around the cranial and caudal ends of the vein. The vein was kept moist using sterile saline and microsurgical scissors were used to make a perpendicular cut between the ligatures through ¾ of the vein width, on the segment most towards the heart where the vein becomes relatively broad. The cranial ligature was then tied with a double knot and 2 pairs of forceps were used to hold the vein in position using the cranial ligature with one hand while inserting the catheter tubing with the other using a dissecting microscope at 10x with a fiber optic light. We used Micro-Renathane® (.037” x .023”) tubing cut at a 60 angle for insertion into the vein. Prior to the surgical procedure, a single length (∼ 1’) of tubing was attached to a 1cc syringe with a 27g needle and loaded with sterile saline. Upon insertion the tubing was secured using the second caudal ligature and the line was evaluated for patency. The tubing was also secured using the cranial ligature and the main incision was closed using surgical nylon.

### 2.3 Intracranial EEG recordings and analysis

Animals were mounted on a stereotaxic frame (David Kopf, Tujunga CA) and an incision was made on the midline of the scalp extending from between the eyes to the back of the skull. The skull was cleaned using gauze and forceps and allowed to dry for several minutes. Bregma was identified and the following coordinates were used to navigate to the BLA: AP -3.0 mm, ML: 5.4 mm, DV: 7.5 mm. A small hole was drilled through the skull using a Dremel tool outfitted with a 1mm dental drill bit. A Teflon-coated stainless-steel wire (#792100, A-M Systems, Sequim, WA) was then inserted into the BLA guided by the stereotaxic instrument and was left in place throughout the remainder of the experiment. EEG data recorded from this wire, relative to an ear clip ground, were low-pass filtered at 100 Hz using a DP-311 differential amplifier (Warner Instruments), passed through a Hum Bug in-line noise eliminator (Hum Bug, Biological Research Technology, Ft Lauderdale, FL), digitized at 1 kHz using a MiniDigi 1B (Axon Instruments, Union City, CA), and then recorded to disk using AxoScope 10.7 (Molecular Devices).

For power analysis the EEG signal was divided into 30 second non-overlapping segments. Total spectral power observed between 0.5 and 55 Hz was calculated using the fast Fourier transform in OriginC (Originlab, Northampton, MA). For event detection in the EEG recordings, we used a high-performance custom method developed in Python 3.12. Specifically, the raw signal was detrended and centered by subtracting a copy of the original signal that had been lowpass filtered using a Butterworth filter with a 0.1 Hz, -3 dB cutoff. This detrended and centered signal was then rectified and lowpass filtered using a Butterworth filter with a 10 Hz, -3 dB cutoff. A threshold event value was then defined for each recording as ∼5 times the mean value observed over the first minute of recording. Event times were identified when the signal crossed the threshold value with an upward slope. Minimum event duration was set at 50 msec in order to detect complex spikes and spindles as single events. After detection event times were binned and reported as events per minute.

### 2.4 Monitoring respiratory rate

A small animal respiration pillow (Small animal instruments Inc., Stony Brook, NY) was placed directly under the sternum and was connected to a custom pneumatic sensor. The pressure waveform was digitized at 1 kHz using a MiniDigi 1B digitizer (Axon Instruments) and recorded with AxoScope 10.7 (Molecular Devices). Breaths were detected in Python 3.12 using a method similar but not identical to that described for event detection in the EEG. Specifically, the respiratory signal was lowpass filtered (Butterworth, 4 Hz -3 dB cutoff) to smooth breaths and minimize influence of heartbeats, then detrended and centered by subtracting an aggressively lowpass filtered copy of the original signal (Butterworth, 0.25 Hz -3 dB cutoff). Breath times were identified when the value of this signal crossed zero with an upward slope. After detection breath times were binned and reported as breaths per minute.

### 2.5 IV Drug Delivery

Following placement of the recording electrode in the BLA, isoflurane was adjusted for delivery of 2% and a brief rest period (5 - 10 min) was used for stabilization of the EEG and respiratory rate. Baseline recordings of 5 min in all experiments were followed by intravenous administration of the OP-based AChE inhibitor NIMP from 5 - 12 minutes. Countermeasures (atropine, HI-6, pirenzepine [PZP], or 2-PAM) were either co-applied (also intravenously) during the last 5 minutes of NIMP exposure (from 7 - 12 min, early intervention), or were applied alone following NIMP (13-17 min, late intervention). A 1cc B&D syringe was used to deliver all drugs. Stock concentrations of all compounds applied intravenously were diluted to final concentrations with sterile saline and infused at a rate of 0.1 cc/min. Total volumes ranged from 0.4 – 0.7 cc. Recordings were continued for at least 20 min after the baseline period when NIMP was delivered at 0.5 mg/kg, and for up to 55 min when NIMP was delivered at 0.375 mg/kg. At the conclusion of recordings animals were heavily anesthetized with 5% isoflurane for > 5 min, removed from the experimental equipment and humanely euthanized.

### 2.6 Statistical analyses

EEG and respiratory data are illustrated normalized to the baseline mean, as obtained during the first 5 minutes of recording. Within groups, a paired Student’s t-test was used to evaluate the effect NIMP (10 – 14 minutes) and countermeasures (19 - 21 minute) relative to the baseline mean (0 - 5 minutes. Across groups, an unpaired Student’s t-test was used to compare data at matching time points. Differences were considered significant where *p* ≤ 0.05.

## 3 Results

### 3.1 Recording BLA EEG and respiratory rate under isoflurane anesthesia

Under isoflurane anesthesia, BLA baseline EEG recordings displayed similarities to cortical sleep patterns. Background voltage oscillations were accompanied by intermittent discharges, often forming complex spindle events with peak-to-peak amplitudes of ∼50 - 150 µV (Figure 2A baseline). As expected, both respiratory rate and EEG activity showed a close correlation with the delivered concentration of isoflurane (data not shown). Recognizing and considering this correlation is crucial when utilizing the current assay. We found that high isoflurane concentrations were not suitable for our studies as steady respiration rates were unattainable, and breathing became irregular and shallow after a 5-minute increase from 2 to 4% isoflurane (not illustrated). However, during constant delivery of 2% isoflurane the baseline EEG recordings were similar across animals and respiratory rates were reliably between 42 - 59 BPM. This facilitated clear detection of CNS hyperactivity and respiratory depression following NIMP infusion and allowed assessment of the efficacy of conventional and novel MCMs. For this reason, 2% isoflurane was used throughout the entire time course of all experiments.

**Figure 2:**
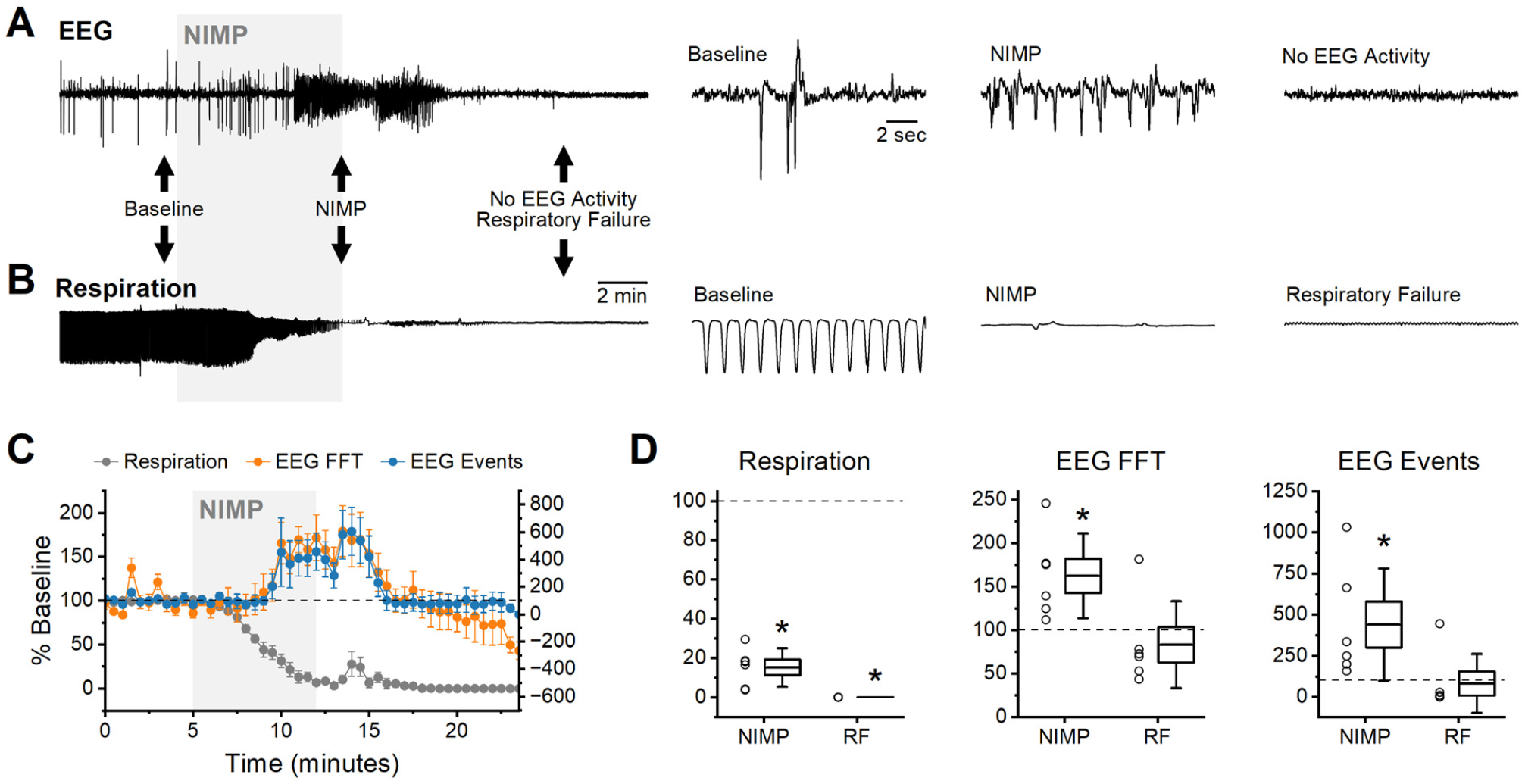
Intravenous NIMP infusion consistently produced seizure-like EEG activity in the BLA and respiratory depression. **(A)** Raw data showing BLA EEG voltage traces before (baseline), during NIMP infusion (NIMP), and after infusion. Left, a 25 min recording. Right, segments from indicated times showing shorter time courses as indicated by black arrows on left. **(B)** Raw data showing individual breaths before (baseline), during NIMP infusion (NIMP), and after respiratory failure (RF). **(C)** Time series data from animals receiving 0.5 mg/kg NIMP. Values are expressed as a percent of baseline + SEM (0 - 5 min) showing respiration rate (gray points, left Y axis), EEG spectral power (orange points, left Y axis), and EEG event frequency (blue points, right Y axis). **(D)** Box plots illustrating effects observed at 10 - 14 min (NIMP) and during respiratory failure (RF, 19 - 21min) in each animal tested. Respiratory rate (left), EEG analysis with FFTs (middle), and EEG event frequency (right). For each box plot, whiskers represent the SD, box ends represent the SE, and the horizontal line inside the box represents the group mean. Asterisks indicate mean difference relative to baseline with p < 0.05 (see Table 1 for further details).

### 3.2 NIMP-induced BLA EEG hyperactivity and respiratory depression

Seizure-like EEG activity in the BLA and respiratory depression were reliably and readily apparent under 2% isoflurane immediately following a five-minute intravenous infusion of 0.5 mg/kg of NIMP without MCMs (Figure 2 A&B). Subsequent analyses revealed a significant reduction in respiratory rate, which was accompanied by a significant increase in both spectral power and event frequency as observed in the EEG (0 - 5 min vs. 10 - 14 min, Figure 2C, Table 1). After an additional 5 minutes, animals were in complete respiratory failure, and EEG activity decreased substantially, likely due to hypoxia (19 - 21 min, Figure 2C & D, Table 1). Additional experiments evaluated the effect of intravenous delivery of both muscarinic antagonists and AChE reactivators as MCMs on these outcome measures. Effects of MCMs were assessed when delivered either during or after acute exposure to NIMP.

**Table 1:**
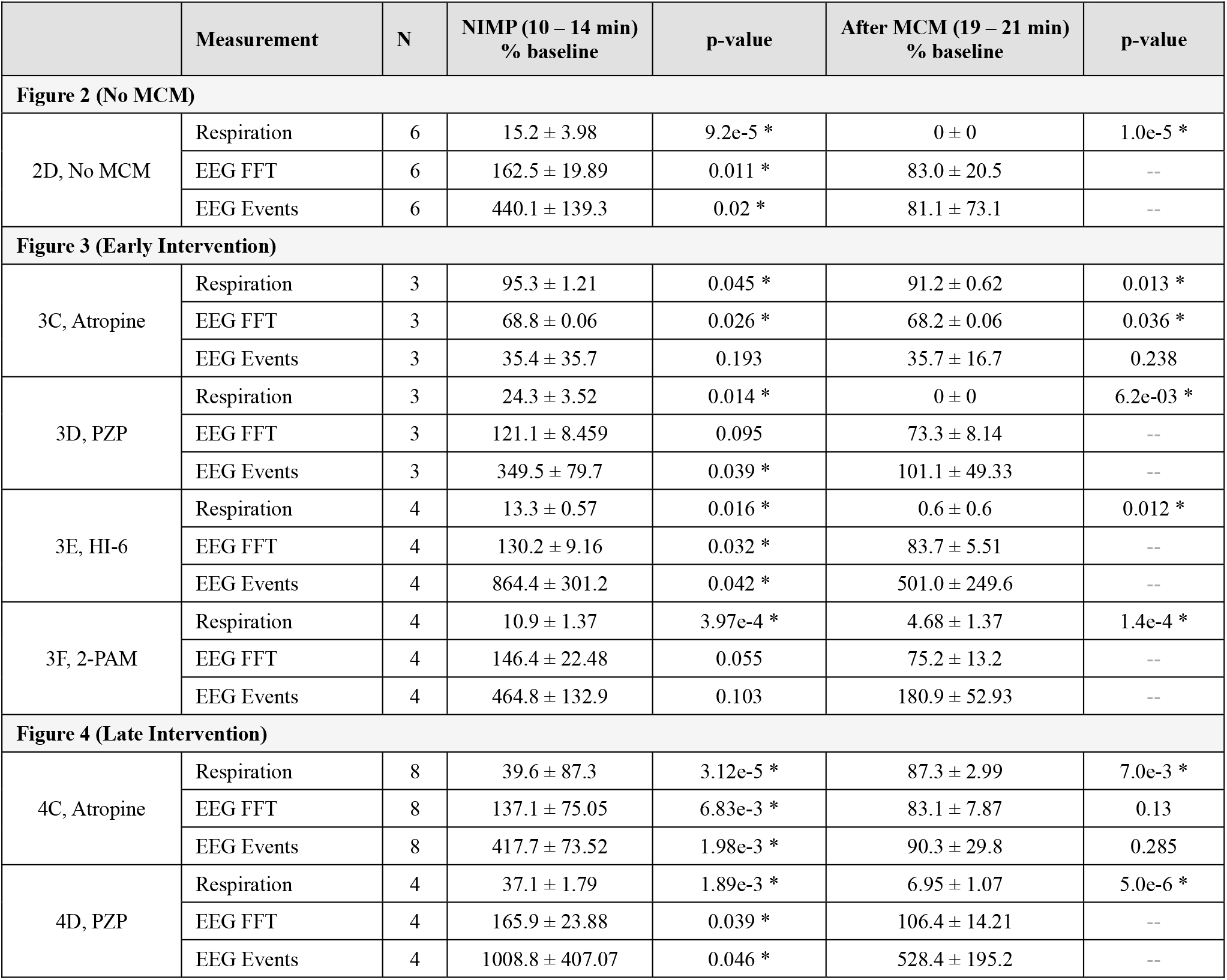
Effects of IV infusion of NIMP and medical countermeasures (MCMs). Respiration (breaths/minute), EEG FFT power (mV^2^/Hz), and EEG event frequency (events/minute) are presented in the indicated time period normalized to the baseline mean (as observed from 0 – 5 minutes). Paired t-tests were run on raw data to compare each of the test periods to baseline. Asterisks indicate comparisons where p < 0.05. For the MCM test period (19-21 minutes) p-values are not reported in cases where respiratory rate is < 10% of baseline, as hypoxia induced by respiratory failure (rather than MCMs) is likely contributing to reduced spectral power and event frequency in the EEG in those cases. Note that in Fig. 3C, EEG spectral power is reduced from baseline during the countermeasure test period suggesting that early intervention with atropine not only prevents NIMP mediated excitotoxicity in the CNS, but likely also reduces cholinergic signaling below basal levels.

|  | Measurement | N | NIMP (10 – 14 min)<br>% baseline | p-value | After MCM (19 – 21 min)<br>% baseline | p-value |
| --- | --- | --- | --- | --- | --- | --- |
| **Figure 2 (No MCM)** |  |  |  |  |  |  |
| 2D, No MCM | Respiration | 6 | 15.2 ± 3.98 | 9.2e-5 * | 0 ± 0 | 1.0e-5 * |
|  | EEG FFT | 6 | 162.5 ± 19.89 | 0.011 * | 83.0 ± 20.5 | -- |
|  | EEG Events | 6 | 440.1 ± 139.3 | 0.02 * | 81.1 ± 73.1 | -- |
| **Figure 3 (Early Intervention)** |  |  |  |  |  |  |
| 3C, Atropine | Respiration | 3 | 95.3 ± 1.21 | 0.045 * | 91.2 ± 0.62 | 0.013 * |
|  | EEG FFT | 3 | 68.8 ± 0.06 | 0.026 * | 68.2 ± 0.06 | 0.036 * |
|  | EEG Events | 3 | 35.4 ± 35.7 | 0.193 | 35.7 ± 16.7 | 0.238 |
| 3D, PZP | Respiration | 3 | 24.3 ± 3.52 | 0.014 * | 0 ± 0 | 6.2e-03 * |
|  | EEG FFT | 3 | 121.1 ± 8.459 | 0.095 | 73.3 ± 8.14 | -- |
|  | EEG Events | 3 | 349.5 ± 79.7 | 0.039 * | 101.1 ± 49.33 | -- |
| 3E, HI-6 | Respiration | 4 | 13.3 ± 0.57 | 0.016 * | 0.6 ± 0.6 | 0.012 * |
|  | EEG FFT | 4 | 130.2 ± 9.16 | 0.032 * | 83.7 ± 5.51 | -- |
|  | EEG Events | 4 | 864.4 ± 301.2 | 0.042 * | 501.0 ± 249.6 | -- |
| 3F, 2-PAM | Respiration | 4 | 10.9 ± 1.37 | 3.97e-4 * | 4.68 ± 1.37 | 1.4e-4 * |
|  | EEG FFT | 4 | 146.4 ± 22.48 | 0.055 | 75.2 ± 13.2 | -- |
|  | EEG Events | 4 | 464.8 ± 132.9 | 0.103 | 180.9 ± 52.93 | -- |
| **Figure 4 (Late Intervention)** |  |  |  |  |  |  |
| 4C, Atropine | Respiration | 8 | 39.6 ± 87.3 | 3.12e-5 * | 87.3 ± 2.99 | 7.0e-3 * |
|  | EEG FFT | 8 | 137.1 ± 75.05 | 6.83e-3 * | 83.1 ± 7.87 | 0.13 |
|  | EEG Events | 8 | 417.7 ± 73.52 | 1.98e-3 * | 90.3 ± 29.8 | 0.285 |
| 4D, PZP | Respiration | 4 | 37.1 ± 1.79 | 1.89e-3 * | 6.95 ± 1.07 | 5.0e-6 * |
|  | EEG FFT | 4 | 165.9 ± 23.88 | 0.039 * | 106.4 ± 14.21 | -- |
|  | EEG Events | 4 | 1008.8 ± 407.07 | 0.046 * | 528.4 ± 195.2 | -- |

### 3.3 Early Intervention with MCMs against NIMP-induced EEG hyperactivity and respiratory depression

Two distinct experimental protocols were implemented to evaluate the capacity to prevent or reverse seizure-like activity and respiratory depression induced by NIMP infusion. These protocols were designated as “early intervention” and “late intervention.” In early intervention experiments, a 5-minute baseline period was followed by NIMP infusion from 5 to 12 minutes. Countermeasures were co-infused with NIMP from 7 to 12 minutes (Figure 3).

**Figure 3:**
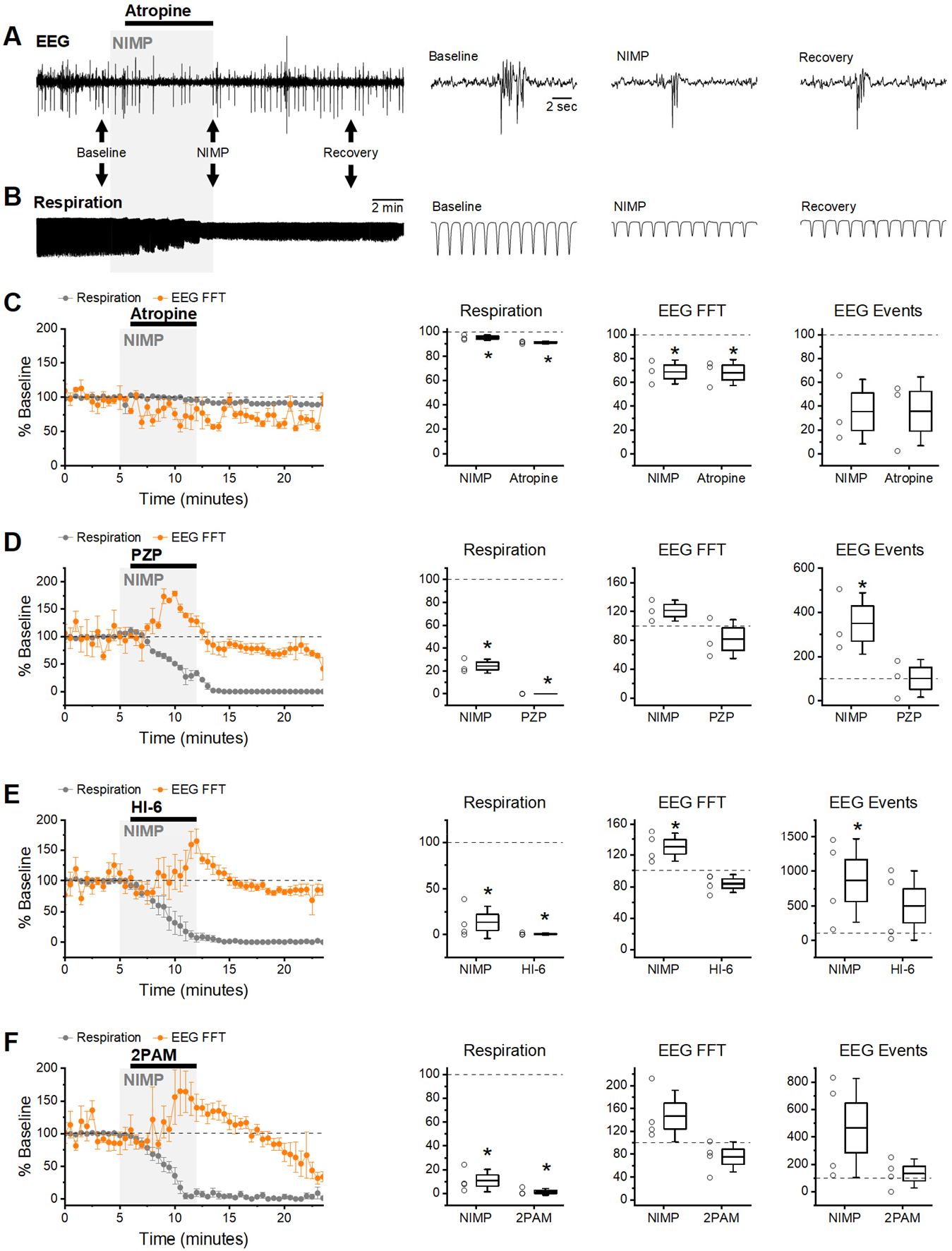
Early Intervention: infusion of MCMs starting 2 minutes after exposure to NIMP. **(A)** Raw data from an experiment using atropine (5 mg/kg) as an MCM. BLA EEG voltage traces before (baseline), during NIMP and atropine infusion (NIMP), and after infusion (Recovery). Left, a 25 min recording. Right, segments from indicated times showing shorter time courses as indicated by black arrows on left. **(B)** Raw data showing individual breaths before (baseline), during NIMP and atropine infusion (NIMP), and after (Recovery) infusion. **(C - F) Left**, group data time series plot from animals receiving 0.5mg/kg NIMP and various MCMs as indicated. Values are expressed as a percent of baseline + SEM (0 – 5min) showing respiration rate (gray points) and EEG data analyzed with FFTs (orange points). **(C - F) Right**, box plots illustrating effects observed after infusion of NIMP (10 -14 min) and countermeasures (19 -21 min) in each animal tested. Respiratory rate (left), EEG spectral power (middle), and EEG event frequency (right). For each box plot, whiskers represent the SD, box ends represent the SE, and the horizontal line inside the box represents the group mean. Asterisks indicate mean difference relative to baseline with p < 0.05 (see Table 1 for further details).

In all instances, EEG seizure-like activity and respiratory depression following NIMP infusion was successfully averted by early intervention with 5 mg/kg atropine (Figure 3 A&B). Indeed, spectral power was reduced relative to baseline following early intervention with atropine, likely due to inhibition of basal cholinergic signaling, while respiratory rate was not significantly different than baseline (Figure 3C, Table 1).

In contrast, early intervention with PZP (5 mg/kg) was ineffective for preventing NIMP induced increases in EEG spectral power and respiratory depression (Figure 3D). Spectral power and event frequency were increased following NIMP infusion and early intervention with PZP, and respiratory depression leading to respiratory failure remained evident, ultimately resulting in a significant decrease in the EEG spectral power and event frequency (Figure 3D, Table 1).

HI-6 (20 mg/kg) and 2-PAM (80 mg/kg) were also ineffective in preventing NIMP-induced EEG seizure-like activity and respiratory depression in early intervention experiments (Figure 3E & 3F, Table 1). Respiratory failure occurred in all animals, ultimately resulting in a decline in event frequency and spectral power of the EEG following infusion of NIMP with HI-6 or 2-PAM (Figure 3E-F, Table 1).

### 3.4 Late intervention with MCMs against NIMP-induced EEG hyperactivity and respiratory depression

For late intervention experiments, a 5-minute baseline period was followed by NIMP infusion from 5 to 12 minutes, and countermeasures were applied from 13 to 17 minutes (Figure 4).

**Figure 4:**
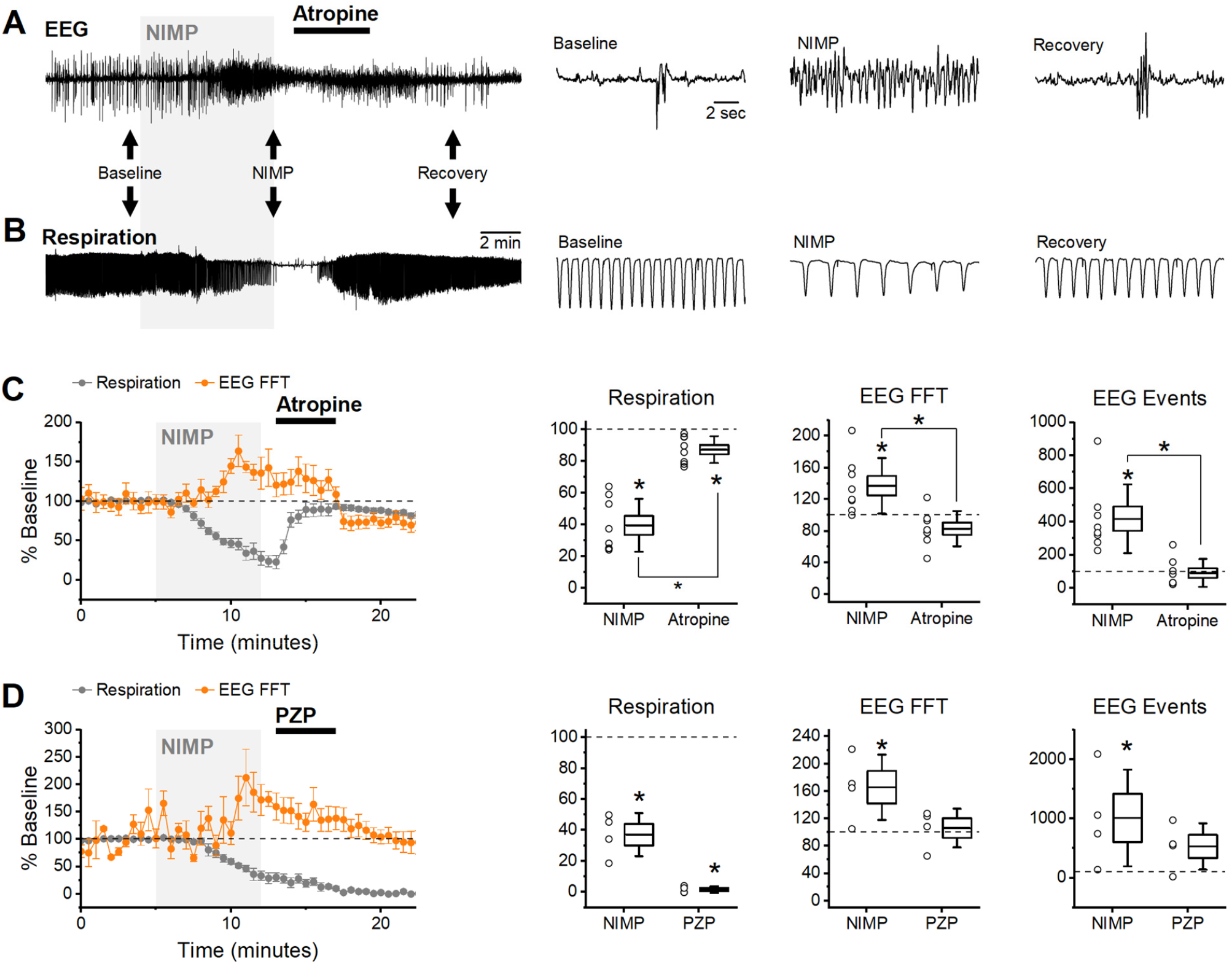
Late Intervention: infusion of MCMs starting 8 minutes after exposure to NIMP. **(A)** Raw data from an experiment using atropine (5mg/kg) as an MCM. BLA EEG voltage traces before (baseline), during NIMP and atropine infusion (NIMP), and after infusion (Recovery). Left, a 25 min recording. Right, segments from indicated times showing shorter time courses as indicated by black arrows on left. **(B)** Raw data showing individual breaths before (baseline), during NIMP and atropine infusion (NIMP), and after (Recovery) infusion. **(C & D) Left**, group data time series plot from animals receiving 0.5 mg/kg NIMP and 5mg/kg atropine (C) or NIMP and 5mg/kg PZP. Values are expressed as a percent of baseline + SEM (0 - 5min) showing respiration rate (gray points) and EEG data analyzed with FFTs (orange points). **(C & D) Right**, box plots illustrating effects observed after infusion of NIMP (10 - 14 min) and countermeasures (19 - 21 min) in each animal tested. Respiratory rate (left), EEG spectral power (middle), and EEG event frequency (right). For each box plot, whiskers represent the SD, box ends represent the SE, and the horizontal line inside the box represents the group mean. Asterisks indicate mean difference relative to baseline with p < 0.05 (see Table 1 for further details). Brackets with asterisks indicate a significant difference between data observed at 10 – 14 min vs. 19 – 21 minutes.

In these experiments, infusion of atropine (5 mg/kg) consistently reversed both electrographic seizure-like activity and respiratory rate depression (Figure 4A & B). Initially, NIMP infusion produced a significant increase in EEG spectral power and event frequency, and a reduction in respiratory rate, however, subsequent infusion of atropine reversed all of these effects to near baseline levels (Figure 4C, Table 1).

In contrast, PZP (5 mg/kg) was ineffective at reversing electrographic seizure-like activity and respiratory depression in late intervention experiments (Figure 4D). Initially, NIMP infusion produced a significant reduction in respiratory rate and increased EEG spectral power and event frequency (Figure 4D, Table 1). Following PZP infusion, respiratory rate did not recover. While spectral power and EEG event frequency did gradually decline from peaks observed during NIMP infusion, this was likely due to hypoxia produced by progressive respiratory failure (Figure 4D, Table 1). Based on these data, and their lack of efficacy in early intervention experiments, HI-6 and 2-PAM were not tested in late intervention experiments.

Overall, data obtained in early and late intervention experiments demonstrate a powerful *in vivo* assay that may be used to screen novel compounds for their ability to prevent or reverse both central and peripheral effects of acute OP-poisoning. While not quite as high throughput as the *in vitro* assay we previously developed, the methods described here represent an important further step in preclinical lead optimization in that they provide a clear way to quantify peripheral vs. central effects of OPs in an anesthetized animal model, and readily indicate whether candidate countermeasures have low vs. high permeability to the BBB.

### 3.5 Sub-lethal doses of NIMP have minimal effect on intracranial EEG but allow evaluation of MCM effects on respiratory rate

In experiments presented above, intravenous delivery of 0.5 mg/kg NIMP, absent effective countermeasures, consistently elicited seizure-like EEG activity and respiratory depression followed by respiratory failure. By contrast, we found that intravenous infusion of 0.375 mg/kg NIMP led to respiratory depression without producing substantial seizure-like EEG activity and did not progress to full respiratory failure. This observation indicated an opportunity to test the effects of MCMs with limited BBB permeability on respiratory rate in live animals over a longer period of time. Toward that end, we tested HI-6 using the late intervention protocol described above, with the lower dose of NIMP, and monitored respiratory rate out to 60 minutes. Results from these experiments revealed that HI-6 infusion was effective in reducing overall respiratory depression produced by infusions of 0.375 mg/kg NIMP. Specifically, this lower dose of NIMP reduced respiratory rate to 47.8 ± 4.9% of baseline in animals that received no countermeasures, and to 68.3 ± 8.1% in animals receiving NIMP followed by HI-6 (Figure 5, p < 0.05). Overall, these results reveal an additional method to evaluate efficacy of novel reactivators as MCMs against acute but sub-lethal effects resulting from exposure to OPs.

**Figure 5:**
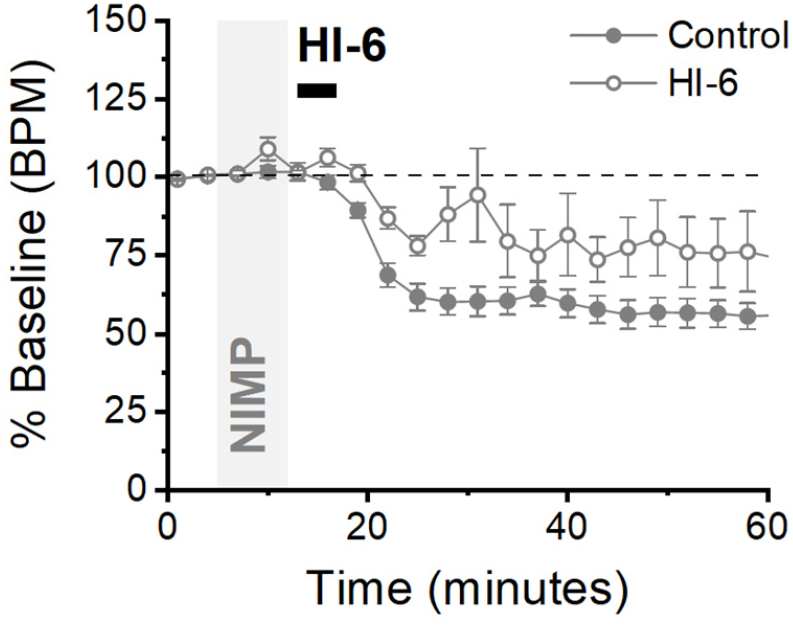
Lower doses of NIMP produce mild respiratory depression that is partially rectified following HI-6 infusion. Time series plot from animals receiving 0.375 mg/kg NIMP (filled circles) and animals receiving NIMP + HI-6 (open circles). Normalized data showing respiratory depression relative to baseline (0 - 5 min) expressed as the mean + SEM for each group over 60 min. Control (n = 7), HI-6 (n = 4).

## 4 Discussion

In the present study conventional MCMs were used to validate a novel whole-animal assay that enabled assessment of MCMs against OP poisoning. The data presented are intended to illustrate a method for evaluating the effectiveness of novel mAChR antagonists and AChE reactivators. The assay allows consistent detection of OP-induced EEG seizure-like activity along with respiratory depression during acute exposure to OPs and during delivery of countermeasures. Subsequent EEG spectral power and event detection analyses enabled statistical comparisons that evaluated the effects of NIMP delivered with and without countermeasures. Our findings aligned with the known ability of specific MCMs to cross the BBB. For example, AChE reactivators and PZP exhibit limited permeability and were ineffective against OP-induced effects, while atropine readily crosses the BBB and was effective at preventing and reversing seizure-like activity and respiratory depression.

Comparing datasets obtained from brain slice experiments to those derived from animals under isoflurane anesthesia yields distinctive insights. The relative efficacy of MCMs in reversing OP effects in brain slices vs. after systemic administration serves as an indicator of their ability to permeate the BBB. Specifically, instances where MCMs exhibit effectiveness in brain slices but prove ineffective when administered systemically suggest a limitation in crossing the BBB. This trend is exemplified by the outcomes associated with HI-6 and PZP, agents with limited BBB penetration, demonstrating inefficacy following systemic administration but proving effective in counteracting NIMP-induced hyperexcitation in brain slices [18]. Additional factors may have played a role in the limited effectiveness of AChE reactivators as MCMs against CNS hyperactivity in the current study. For example, the existing standard of care mandates the administration of these agents alongside atropine, particularly due to OP-induced hyperstimulation of cholinergic receptors which can lead to sustained excitatory effects mediated by other neurotransmitters and refractory to treatment with reactivators. Also, in some cases reactivators may have limited efficacy against irreversible bonds formed between OPs and AChE. In consideration of the inefficacy of reactivators in early intervention experiments (Figure 3E & 3F), we confirmed the ability of HI-6 to partially reverse non-lethal respiratory depression produced by lower doses of NIMP (Figure 5). Overall, the data presented here underscore the utility of combining the current assay with brain slice studies [e.g. see 18] to assess BBB permeability and enable evaluations of novel MCMs regarding their ability to mitigate OP-induced CNS effects.

EEG recordings were evaluated using power spectral analysis and by using threshold-based event detection software designed to detect complex spikes in the BLA (Figure 2A, baseline). From both a qualitative and quantitative perspective, both methods produced very similar results. That said, spectral analysis allows effects to be quantified in specific / narrow frequency bands, and this may provide additional utility in some cases.

It should be noted that while employing these analytical tools it is imperative to consider the apparent reversal of EEG hyperactivity in the absence of MCM infusion or following ineffective MCM treatments. Specifically, respiratory failure reverses NIMP induced increases in spectral power, likely due to hypoxia. Thus, reversal of NIMP induced increases in spectral power should not be considered indicative of an effective MCM treatment unless respiratory rate also recovers.

The results of our studies necessarily involve the interplay between isoflurane, neurotransmitters, and OPs. Therefore, it is crucial to acknowledge the effects of isoflurane on glutamatergic neurotransmission. The anesthetic properties of isoflurane are likely attributable to its impact involving the inhibition of glutamate release, downregulation of NMDA receptor expression, reduction of EPSC frequency and amplitude, and increase in GABA _A_ receptor-mediated responses [25–27]. However, infusion of NIMP at 0.5 mg/kg consistently induced seizure-like BLA EEG activity in animals anesthetized with 1 - 3% isoflurane, a concentration range which has been recommended for full surgical plane maintenance [28]. In essence, despite isoflurane’s broad effects on multiple neurotransmitter systems, including glutamate, the residual EEG activity during anesthesia was sufficient to detect OP-induced CNS hyperactivity. This aligns with research demonstrating that the AMPA receptor antagonist NBQX, while effectively mitigating neuronal toxicity in rodents exposed to OPs, does not completely eliminate OP-induced increases in CNS excitability OP-induced seizures when administered alone [29]. In a related context, recent findings by Puthillathu et al. [30] show that brief exposure to high concentrations of isoflurane (up to 5%) delivered 30 - 180 minutes following OP exposure effectively controls convulsions and mitigates neuropathology, presumably by dampening glutamatergic toxicity. However, our assay focuses on therapeutic interventions administered within 2-8 minutes following the onset of OP infusion, and its relevance lies in predicting the ability to prevent or reverse CNS seizure-like EEG activity well before the long-term effects of neurotoxicity manifest. In this regard, our model appears most pertinent for scenarios involving self-administration or timely first responder intervention.

Due to the propensity of certain OPs to induce persistent convulsions, established medical countermeasure protocols involve the administration of benzodiazepines [31]. Notably, in conscious animals, NIMP doses comparable to those employed in our study typically induce convulsions and other seizure-like behaviors [32], which in our model are blocked by general anesthesia. Consistent with this are results showing isoflurane demonstrates potent anticonvulsant and neuroprotective efficacy in a paraoxon model of OP poisoning [33]. Nonetheless, the rectification of EEG seizure-like activity under isoflurane may serve as a sufficient and viable indicator for predicting therapeutic outcomes in both conscious animal models and humans. Importantly, this insight may enable experimenters to avoid subjecting animals to the detrimental consequences associated with cholinergic crises.

Isoflurane is a common choice for small animal surgeries. It offers numerous advantages compared to other anesthesia types, such as swift induction and recovery, stability, precision, cardiovascular and respiratory stability, and a wide safety margin. The key benefit of using isoflurane in the current assay is that it allows the experimenter to maintain stable basal EEG activity over an extended period of time. This allows the isolation of neuronal hyperexcitation induced by OPs and the evaluation of MCMs without concerns about confounding effects related to fluctuations in anesthesia depth. Ketamine and xylazine combinations are also prevalent in laboratories conducting rodent surgeries. They work in tandem to provide analgesia through their actions on NMDA receptors and α2 adrenergic receptors [34]. Additionally, urethane finds widespread use in electrophysiological animal studies due to its minimal impact on cardiovascular and respiratory systems and its preservation of spinal reflexes. Recent studies indicate its effects on multiple neurotransmitter-gated ion channels, including GABA, NMDA, AMPA, and nicotinic receptors [35]. The most common administration route for both ketamine/xylazine and urethane is intraperitoneal (IP) injection. While stable EEG recordings may be achieved with proper dose titration and short experiment durations, constant metabolic degradation of these agents occurs, making their use potentially less desirable than continuous delivery of vaporized isoflurane, however see [36]. Moreover, individual responses to these agents can be highly variable, with some animals not reaching an acceptable surgical anesthetic plane [34]. Such variability could confound results obtained using the current assay. Nevertheless, the exploration of alternative anesthetics could be a subject for future investigations, although it was beyond the scope of the current work.

Overall, we believe the methods described here, in combination with an *in vitro* assay previously described [18], provide a robust platform for evaluating the effectiveness of novel AChE reactivators, muscarinic antagonists, and other compounds against both central and peripheral effects of OP poisoning. Future refinements to the current assay could further explore BBB permeability and efficacy in the CNS by comparing the effects of the same compound delivered intravenously vs. by intracerebroventricular injection. Using a respirator in experiments that involve higher doses of NIMP may also allow evaluation of the ability of MCMs to prevent or reverse CNS effects in response to doses of OPs which would otherwise cause lethal respiratory depression. Finally, it is important to highlight that OP-induced cholinergic crises in conscious animals produce overt signs of suffering including chewing, gnawing, profuse secretions, diarrhea, muscle fasciculation, restlessness, tremors, convulsions, and fatal respiratory distress. Implementation of the assay outlined here and studies using brain slices offers a more humane alternative for MCM testing involving conscious animals. As such, this work aligns well with the principle of “refinement” advocated by the international organization AAALAC which promotes humane treatment of animals in science.

## 5. Data Availability Statement

The raw data supporting the conclusions of this article will be made available by the corresponding author, without undue reservation.

## 6. Conflict of Interest

The authors declare that the research was conducted in the absence of any commercial or financial relationships that could be construed as potential conflicts of interest.

## 7. Author Contributions

JST developed the assay, performed experiments, completed data analysis, and wrote the initial version of the manuscript. SH assisted with data analysis, assay development, figure preparation, software development, and manuscript editing. JDT assisted with direction of the project and manuscript editing. CJF assisted with direction of the project, assay development, manuscript writing, production of final figures, and development of software tools for data analysis.

## 8. Funding

Effort sponsored by the U.S. Government under Other Transaction number W15QKN-16-9-1002 between the MCDC, and the Government. The US Government is authorized to reproduce and distribute reprints for Governmental purposes notwithstanding any copyright notation thereon.

## 9 Acknowledgements

We thank Dr. Michael A. King for helpful discussion on experimental design, and Dr. Becky Hood for acquisition, storage, and aliquoting of NIMP, HI-6, and 2-PAM.

## 10 Contribution to the Field Statement

Organophosphates (OPs) act both peripherally and centrally to produce respiratory depression and seizures that often lead to death. Here we describe the development of a novel isoflurane-anesthetized whole-animal model that allows for the recording of sustained OP-induced respiratory depression and electrographic seizure-like activity following intravenous OP delivery. We suggest the assay described can be a valuable tool for preclinical evaluation of novel compounds as medical countermeasures (MCMs) for toxicity produced by acute exposure to OPs. The assay described provides methods for reliably and precisely quantifying both peripheral and central effects of OPs, and the ability of novel MCMs to prevent or reverse such effects. In combination with a previously described *in vitro* assay, current methods also provide information on whether candidate MCMs have low vs. high permeability to the blood brain barrier. The fact that the current study is conducted entirely under general anesthesia, and yet is still able to provide clear and quantitative measures of the effects of OPs and MCMs in the CNS means that it also represents a humane alternative to testing the effects of OPs and MCMs in conscious animals.

## Notes

### Competing Interest Statement

The authors have declared no competing interest.

